# Clinical implications of identifying pathogenic variants in aortic dissection patients with whole exome sequencing

**DOI:** 10.1101/497917

**Authors:** Brooke N. Wolford, Whitney E. Hornsby

## Abstract

**Background:** Thoracic aortic dissection is an emergent life-threatening condition. Routine screening for genetic variants causing thoracic aortic dissection is not currently performed for patients or their family members.

**Methods:** We performed whole exome sequencing of 240 patients with thoracic aortic dissection (n=235) or rupture (n=5) and 258 controls matched for age, sex, and ancestry. Blinded to case-control status, we annotated variants in 11 genes for pathogenicity.

**Results:** Twenty-four pathogenic variants in 6 genes (*COL3A1, FBN1, LOX, PRKG1, SMAD3, TGFBR2*) were identified in 26 individuals, representing 10.8% of aortic cases and 0% of controls. Among dissection cases, we compared those with pathogenic variants to those without and found that pathogenic variant carriers had significantly earlier onset of dissection (41 vs. 57 years), higher rates of root aneurysm (54% vs. 30%), less hypertension (15% vs. 57%), lower rates of smoking (19% vs. 45%), and greater incidence of aortic disease in family members. Multivariable logistic regression showed significant risk factors associated with pathogenic variants are age <50 [odds ratio (OR) = 5.5; 95% CI: 1.6-19.7], no history of hypertension (OR=5.6; 95% CI: 1.4-22.3) and family history of aortic disease (mother: OR=5.7; 95% CI: 1.4-22.3, siblings: OR=5.1; 95% CI 1.1-23.9, children: OR=6.0; 95% CI: 1.4-26.7).

**Conclusions:** Clinical genetic testing of known hereditary thoracic aortic dissection genes should be considered in patients with aortic dissection, followed by cascade screening of family members, especially in patients with age-of-onset of aortic dissection <50 years old, family history of aortic disease, and no history of hypertension.

## INTRODUCTION

Thoracic aortic dissection is a life-threatening condition, responsible for 15,000 deaths a year in the United States.^1^ Up to 25% of patients presenting with a thoracic aortic aneurysm and dissection have an underlying genetic predisposition,^2, 3^ which can be associated with syndromic features, such as Marfan syndrome or Loeys-Dietz syndrome, or not associated with syndromic features, as with *PRKG1* and *MYH11* mutations.^4^ Variants in many genes, including *FBN1, ACTA2* and *SMAD3*, among others, can lead to either syndromic or non-syndromic thoracic aortic aneurysm and dissection.^5^ Recent advances in the field have shown definitive and strong evidence to support the role of pathogenic variants in *ACTA2, COL3A1, FBN1, MYH11, SMAD3, TGFB2, TGFBR1, TGFBR2, MYLK, LOX*, and *PRKG1*^5^ as predisposing to hereditary thoracic aortic disease.^3^

These genetic findings play a critical role for the patient and family members, helping to guide clinical decision-making to prevent or lessen the impact of a life-threatening thoracic aortic event. Aortic diameter is a central criterion when deciding prophylactic surgical intervention, and the recommended aortic diameter for surgical intervention differs for those with and without an underlying genetic predisposition. The American Heart Association/American College of Cardiology (AHA/ACC) guidelines^6^ recommend that patients with genetically mediated aneurysms undergo elective surgical repair at an ascending aortic diameter of 4.0-5.0 cm, depending on the condition; whereas patients without a known genetic mutation undergo elective surgical repair when the sinus or ascending aortic diameter is ≥ 5.5 cm. Recent work shows that different genes predisposing to hereditary thoracic aortic dissection have varying presentations and courses.^7, 8^ For instance, patients with *ACTA2* mutations more often present with acute aortic dissections whereas patients with Marfan syndrome often present with skeletal and ocular features before aortic dilation is discovered.^9^

When a patient presents with a thoracic aortic dissection, it is commonly unknown whether the patient has an underlying rare genetic variant triggering the dissection. The identification of variants known to predispose to thoracic aortic dissection has the potential to improve clinical management and guide treatment strategies for patients and family members. The objective of this study is to examine the utility of research-level whole exome sequencing in patients with a history of thoracic aortic dissection or rupture, as well as to identify which patients and corresponding family members may be most likely to benefit from clinic genetic testing.

## METHODS

### Study Design

The Cardiovascular Health Improvement Project (CHIP) is a biorepository with a historical collection of phenotype data, family history, DNA, genotypes, and aortic tissue samples from participants with aortic disease. Between August 2013 and December 2015, 1752 participants were enrolled in the CHIP biorepository, and of those, 265 cases had a diagnosis of thoracic aortic dissection including type A or type B aortic dissection or aortic rupture with or without aortic aneurysm. Age-, sex-, and ancestry-matched controls (n=265) were identified as previously described from the Michigan Genomics Initiative (MGI), which is a surgical-based biobank.^10^ In brief, we matched thoracic aortic dissection cases from CHIP to MGI controls of the same sex, age range (−5, +10) at time of enrollment, and minimum Euclidean distance as calculated from the first two principal components of genotype data indicative of genetic ancestry. Principal components were obtained by principal component analysis (PCA) in PLINK 1.9^11^ (www.cog-genomics.org/plink/1.9) on 58,563 genotyped variants with > 0.05 minor allele frequency. For 83 CHIP samples without genotypes from a customized Illumina HumanCoreExome v12.1 bead array, we used self-reported ancestry instead of principal component-based ancestry to identify controls with similar genetic ancestry. In the event that insufficient DNA was available for the best matched control, we moved sequentially through the top 10 best matched controls. All study procedures were approved by the Institutional Review Board (HUM00052866 and HUM00094409).

### Clinical Characteristics

For the thoracic aortic dissection cases (hereon referred to as cases) identified from CHIP, the researchers leveraged the electronic medical record (EMR) to verify demographics, clinical diagnoses, family history, surgical history, Clinical Laboratory Improvement Amendments (CLIA) genetic testing results, syndromic features, medications, and comorbidities. We cross-referenced the clinical diagnosis with CLIA and research-level whole exome sequencing to examine continuity between diagnoses. Two researchers performed all EMR interpretations. Clinical characteristics for the cases are presented as median and inter-quartiles for continuous data and n (%) for categorical data. Univariate comparisons were performed using Chi-square with Yates’ continuity correction or Fisher’s exact test when any expected cell counts were < 5 for categorical data, and Wilcoxon rank sum tests were used for continuous data. Multivariable logistic regression was used to identify associations between risk factors and pathogenic variant carriers.

### Whole Blood Processing, DNA Isolation, Plating DNA Samples for Whole Exome Sequencing

DNA from whole blood was isolated using the Gentra Puregene Blood kit (Qiagen) following the manufacturer’s protocol with reagent volumes scaled to whole blood input of 7.5 ml. DNA samples for cases and controls (n=530) were prepared for whole exome sequencing as outlined by the Northwest Genomics Center (NWGC, University of Washington) as follows: (1) plated on 96-well barcoded plates (plates provided by NWGC) with foil plate seals after the plate manifest had been approval, (2) at least 4 micrograms of high-quality genomic DNA was plated per sample, and (3) all DNA samples were normalized to a concentration between 80 to 100 nanograms per microliter (a total volume of at least 50 microliters per sample) and suspended in 1xTE solution (10mM Tris-HCL, pH 8.0, 0.1mM EDTA, pH 8.0) and quantitated using the Quant-iT PicoGreen dsDNA kit (Invitrogen).

### Whole Exome Sequencing

#### Sample Receipt, Quality Control and Tracking

The NWGC centralizes all receipt, tracking and quality control/assurance of DNA samples. Samples have a detailed sample manifest (i.e., identification number/code, sex, DNA concentration, barcode, extraction method). Initial quality control entails DNA quantification, sex-typing and molecular fingerprinting using a high frequency, cosmopolitan genotyping assay. This fingerprint is used to identify potential sample handling errors and provides a unique genetic ID for each sample, which eliminates the possibility of sample assignment errors. Samples are failed if: (1) the total amount, concentration or integrity of DNA is too low; (2) the fingerprint assay produces poor genotype data or (3) sex-typing is inconsistent with the sample manifest. 530 samples were submitted to NWGC for whole exome sequencing and 528 were approved for sequencing with sufficient DNA quantity.

#### Library Production, Exome Capture, Sequencing

Library construction and exome capture have been automated (Perkin-Elmer Janus II) in 96-well plate format. 1 ug of genomic DNA was subjected to a series of shotgun library construction steps, including fragmentation through acoustic sonication (Covaris), end-polishing, A-tailing and ligation of sequencing adaptors with dual 8 bp barcodes for multiplexing, followed by PCR amplification. Libraries underwent exome capture using Roche/Nimblegen SeqCap EZ v2.0 (~36.5 MB target). Prior to sequencing, the library concentration was determined by fluorometric assay and molecular weight distributions verified on the Agilent Bioanalyzer (consistently 150 ± 15 bp).

#### Sequencing

Barcoded exome libraries were pooled using liquid handling robotics prior to clustering (Illumina cBot) and loading. Massively parallel sequencing-by-synthesis with fluorescently labeled, reversibly terminating nucleotides was carried out on the HiSeq sequencer, with 8 exomes multiplexed per lane (60 to 80x mean coverage per sample).

#### Read Processing

NWGC’s sequencing pipeline is a combined suite of Illumina software and other industry standard software packages (i.e., Genome Analysis ToolKit [GATK], Picard, BWA-MEM, SAMTools, and in-house custom scripts) and consisted of base calling, alignment, local realignment, duplicate removal, quality recalibration, data merging, variant detection, genotyping and annotation.

#### Variant Detection

Variant detection and genotyping were performed using the HaplotypeCaller tool from GATK^12^ (3.2). Variant data for each sample was formatted (variant call format [VCF]) as “raw” calls that contain individual genotype data for one or multiple samples. NWGC applied hard filtering on the “raw” VCF to generate a filtered VCF call set (GATK v3.4).

#### Data Analysis Quality Control

Data quality control included an assessment of: (1) total reads (minimum of 40 million PE75 reads); (2) library complexity (3) capture efficiency (4) coverage distribution: 90% at 8X required for completion; (5) capture uniformity; (6) raw error rates; (7) Transition/Transversion ratio (Ti/Tv); (8) distribution of known and novel variants relative to dbSNP (9) fingerprint concordance > 99%; (10) sample homozygosity and heterozygosity and (11) sample contamination validation. Exome completion was defined as having > 90% of the exome target at > 8X coverage and >80% of the exome target at > 20X coverage. Typically this requires mean coverage of the target at 30-60X. A total of 521 samples, 260 cases and 261 controls, and 323,867 variants (single nucleotide polymorphisms and insertion/deletions) passed quality control and were released to researchers.

### Additional sample and variant filtering

Bi-allelic sites were extracted and lower coverage genotypes with depth (DP) < 5 were masked out. All samples met the quality control threshold of an individual level call rate > 0.9. Poor quality sites with site-level call rate < 0.9 were excluded. Variants significantly deviating from HWE with p-value < 1×10^−6^ were also removed. KING^13^ was used to identify five sample pairs as duplicates, and the sample with the lowest call rate was excluded, leaving 258 cases and 258 controls. Concordance with Exome+GWAS array genotypes was > 0.999 across all minor allele frequencies. After review of the medical record nine cases were excluded for traumatic aortic dissection (i.e, accident or illicit drug use) and nine for abdominal aorta rupture as the etiology of disease is typically atherosclerotic in nature. The final analysis set was comprised of 240 cases and 258 controls and 299,195 variants. We opted to keep all cases and controls that passed quality control procedures, rather than reduce the sample size by only including complete pairs.

### Annotation of variants with a clinical impact

We focused on the following genes which confer a dominantly inherited risk for aortic dissection and with definitive and strong evidence of association of hereditary thoracic aortic aneurysm and dissection: *ACTA2, COL3A1, FBN1, MYH11, SMAD3, TGFB2, TGFBR1, TGFBR2, MYLK, LOX*, and *PRKG1*^5,14^. A total of 248 variants in these genes were annotated using dbNSFPv3.5a. and reviewed by a single researcher blinded to case or control status of the sample in which the variant was identified. Variants were then annotated as pathogenic, variants of unknown significance, or benign. Protein isoforms that are major isoforms expressed in smooth muscle cells (SMC) or used in previous publications were used to predict amino acid changes (Supplementary Table 1). To define pathogenic variants, we annotated variants based on the ACMG-AMP standards and guidelines.^15^ Additionally, established rules^14^ were used to classify rare variants as pathogenic or disease-causing. Rare variants were annotated as variants of unknown significance if lacking proof of pathogenicity. Variants were considered benign if they are nonsynonymous mutations with MAF ≥ 0.005 in ExAC Non-Finnish Europeans^16^ or in a nonrelevant isoform, were synonymous mutations, or occurred > ±2 bp from intron/exon boundaries.

### Molecular Inversion Probe Sequencing

Molecular Inversion Probe Sequencing (MIPS) was performed as a technical replicate of cases and controls that were whole exome sequenced and found to carry a pathogenic variant. MIPS was first performed on DNA from the same extraction used for whole exome sequencing. An additional round of MIPS was performed from a second DNA isolation to serve as a sample replicate. A custom targeted sequencing panel was designed for 116 genes using single molecule molecular inversion probes or smMIPS.^17^ Coding exon coordinates were retrieved from the UCSC Genome Browser “knownGene” table (build GRCh37/hg19) and padded by 5 bp in each direction to include splice sites. Probes were designed and prepared as previously described.^18^ Briefly, probe sequences unique to each targeted region were selected using custom Python scripts, and synthesized in batch as a microarray of 150mers (CustomArray, Inc). Probes were designed with identical adaptor sequences and were amplified with PCR using primers directed to those sequences. Adaptors were removed by overnight digestion with EarI (NEB), purified with one volume SPRI beads supplemented with five volumes isopropanol, and eluted in Tris-EDTA pH 8. For each sample, approximately 9 ng of purified smMIPS probes were combined with 250 ng genomic DNA, in a reaction mixture including Ampligase DNA Ligase Buffer 1X (Epicentre), 0.4 uM dNTPs (NEB), 3.2U HemoKlentaq (NEB) and 1U Ampligase (Epicentre). After denaturation at 95°C for 10 minutes and incubation at 60°C for 20 hours, linear probes and the remaining genomic DNA were removed by exonuclease treatment with ExoI and ExoII (NEB). The captured material was amplified by PCR using barcoded primers. The resulting PCR products were pooled for one lane of paired-end 150 bp sequencing on an Illumina HiSeq 4000 instrument at the University of Michigan Sequencing Core. Reads were aligned to the human genome reference (build GRCh37/hg19) using bwa mem^19^ and a custom pipeline (available at https://github.com/kitzmanlab/mimips) was used to remove smMIPS probe arm sequences and remove reads with duplicated molecular tags.

Variant calling of MIPS sequencing results for both single nucleotide variants and insertions/deletions was performed using the GotCloud^20^ pipeline. All sequencing results were aligned to build GRCh37/hg19. An iterative filtering process was performed after variant calling to remove variants with a depth less than 10, then samples with call rates of less than 0.6, followed by variants with a call rate of less than 0.8 and finally samples with call rates below 0.9.

### Statistical analysis for burden of variants in cases and controls

To test for association between carriers of a given variant class and case/control status we used Fisher’s exact test when any expected cell counts were < 5 and Chi-square test with Yates’ continuity correction otherwise. This was done using the statistical programming language R version 3.5.1. We identified first degree relatives using KING2^13^ and whole exome sequencing variant calls. For the two first degree relative pairs we found in the cases, we retained the first sample acquired (proband) for the analysis resulting in 238 cases. A sample carrying at least one of a variant class was considered a carrier. We performed burden tests for association with case status across the 11 genes for all pathogenic variants (N=24) and variants of unknown significance (N=86). We first excluded carriers of pathogenic variants before testing for association with case status for carriers of variants of unknown significance (Ncases=213, Ncontrols=258). A Bonferroni threshold of 0.003 is used to account for 17 independent tests, which are assumed to be independent.

### Data Visualization

Annotated Fibrillin 1 protein domains from Pfam 31.0^21^ were with a modified version of GenVisR 1.14.1^22^ for data visualization. Variants falling in mutation splice sites are not included in this protein-level visualization.

## RESULTS

### Annotation of variants from research-level whole exome sequencing identifies pathogenic variants

A total of 240 cases with a clinical diagnosis of thoracic aortic dissection (type A or type B) or rupture with or without aortic aneurysm and 258 age-, sex-, and ancestry-matched controls had whole exome sequences available following quality control (see Methods) for quality control failures). For the 498 samples passing quality control, 248 variants were annotated blind to the variant carrier’s case or control status. 24 pathogenic variants in 6 genes (*COL3A1, FBN1, LOX, PRKG1, SMAD3, TGFBR2*) were identified, found exclusively in 26 cases (Table 1), representing 10.8% of cases and 0% of controls. Two variants were seen twice in cases who were first degree relatives. There is a significant burden of pathogenic variants in *FBN1* in cases compared to controls (N_cases_=18, N_controls_=0; p-value=2.5×10^−5^, Supplementary Table 2). These variants are predominantly found in calcium-binding epidermal growth factor domains of *FBN1* (Figure 1). We examined the proportion of pathogenic variants that were present in commonly used databases and found that of the 24, 11 were present in dbSNP^23^, 8 were listed as pathogenic in ClinVar^24^, and 2 were present in gnomAD^16^ (see Supplementary Note).

**Figure 1.**
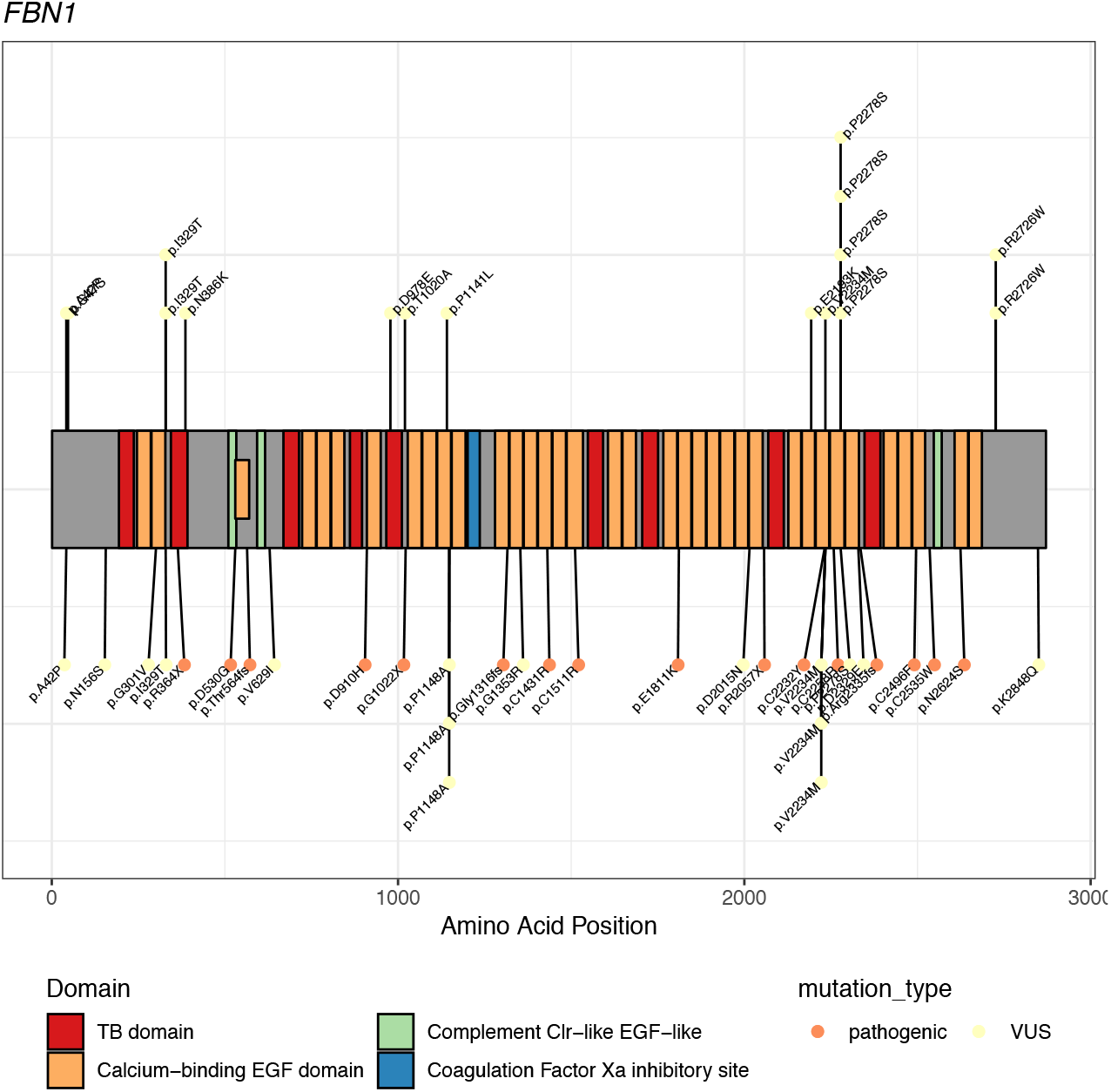
Distribution of pathogenic and variants of unknown significance in Fibrillin 1 (*FBN1*). Each point is a sample, with controls above the protein diagram and cases below.

**Table 1.**
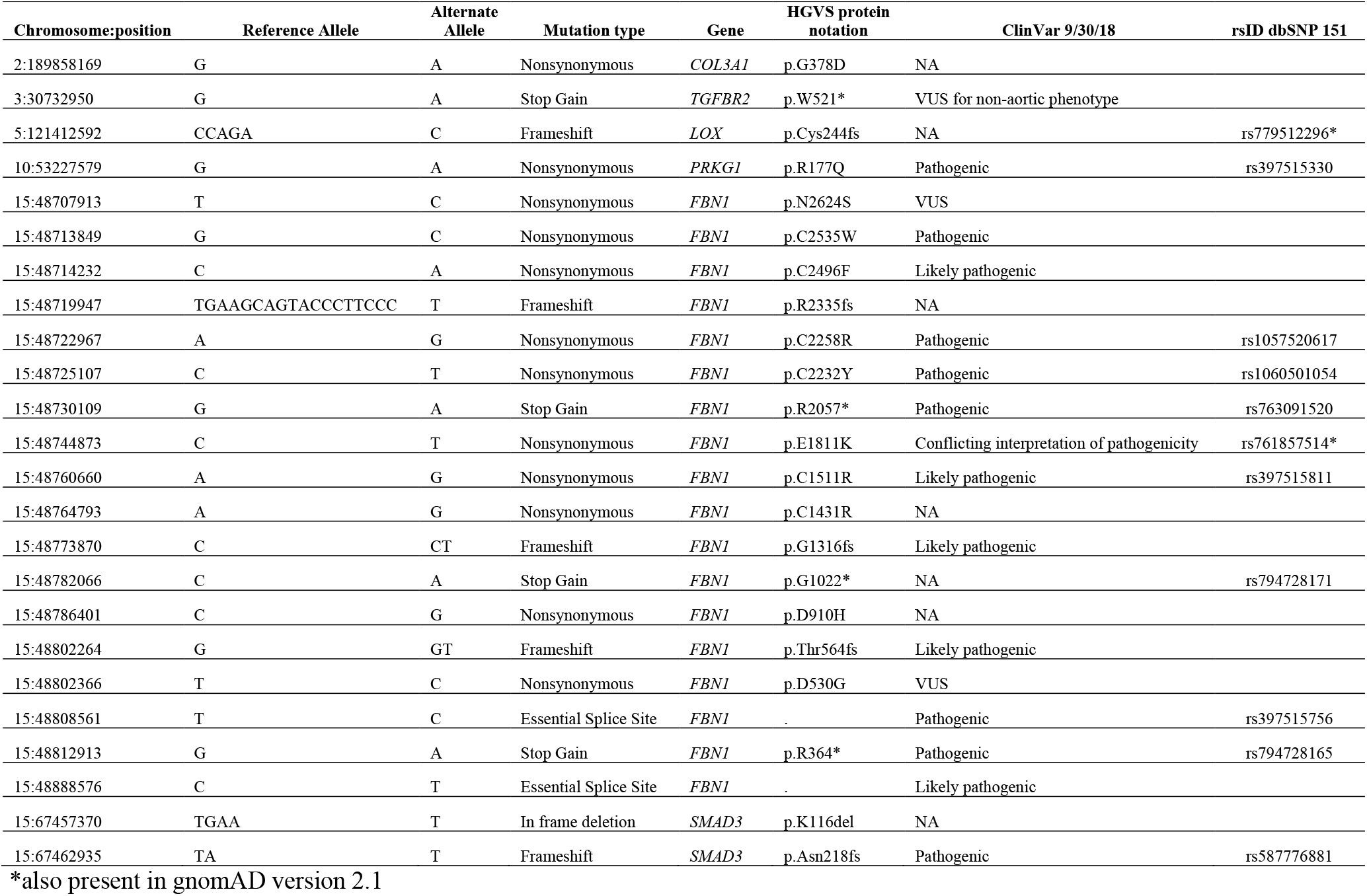
Classification of 24 Pathogenic Variants

### Validation targeted sequencing

Molecular Inversion Probe Sequencing was utilized to ensure the highest level of confidence in the whole exome sequencing variant calls and protect against potential sample swaps. The carrier status of pathogenic variants identified through whole exome sequencing was verified by Molecular Inversion Probe Sequencing in all 26 samples (Supplementary Table 3).

### Research-level whole exome sequencing and precision health

For 17 of the 26 pathogenic variant carriers (hereon pathogenic carriers), the research-level whole exome sequencing results aligned with the current clinical diagnoses in the EMR, including 5 patients (5/17) in which CLIA genetic testing previously identified the same pathogenic variant (Supplementary Table 4). However, for 9 pathogenic carriers, research-level whole exome sequencing and annotation of pathogenic variants added clinical precision to the current clinical diagnosis (Table 2). 8 of these cases (8/9) lacked a specific clinical diagnosis and research-level whole exome sequencing revealed Marfan syndrome *(FBN1*, n=4), vascular Ehlers-Danlos syndrome *(COL3A1*, n=1), or familial thoracic aortic disease *(LOX, PRKG1*, and *SMAD3*, n=3). For 1 pathogenic carrier (1/9) there was an incorrect diagnosis of Marfan syndrome, which was revised to Loeys-Dietz syndrome based on a pathogenic variant identified in *TGFBR2*. In addition, the research-level whole exome sequencing results provide a basis for clinical cascade screening for the family members of all 26 cases (Table 2).

**Table 2.**
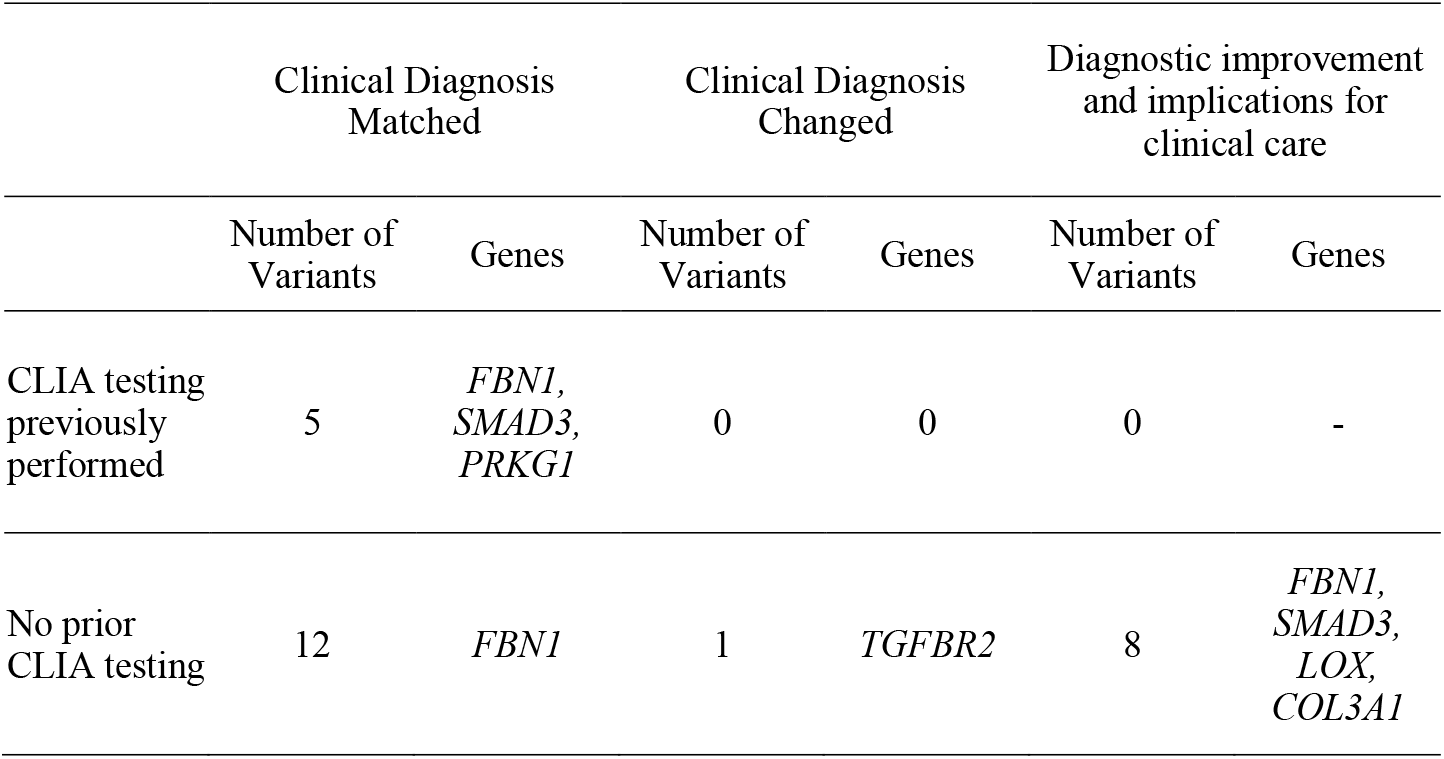
Comparison Between Clinical Diagnosis and Pathogenic Variants

### Variants of unknown significance

86 of the 248 annotated variants in aortopathy genes were annotated as variants of unknown significance. After excluding one of each 1^st^ degree relative pair (see Methods) and cases with pathogenic variants, 58 of 213 cases (27.2%) and 51 of 258 controls (19.8%) have at least one variant of unknown significance identified from whole exome sequencing, which was not significant (p-value=0.072, Supplementary Table 5). There is, however, a significant association between pathogenic variants and cases (p-value=2.8×10^−7^, Supplementary Table 5). None of the 11 genes demonstrated association between carrier status for variants of unknown significance and thoracic aortic dissection or rupture case/control status (Supplementary Table 2).

### Clinical characteristics between pathogenic variant and non-pathogenic variant carriers

The pathogenic carriers were significantly younger (41 vs. 57 years, 77% younger than 50 years old), had significantly more root aneurysms (54% vs. 30%), less hypertension (15% vs. 57%) and less history of smoking (19% vs. 45%) compared to the non-pathogenic carriers. Moreover, the pathogenic carriers had a greater incidence of aortic disease in parents, siblings, and children (all p-values<0.05) (Table 3). Pathogenic carriers presented with more type A than type B dissections although this comparison was not significant (69.2% vs. 58.9%, p=0.421). Multivariable logistic regression showed that pathogenic carriers were significantly more likely to have dissection age < 50 years old, family history of aortic disease, and no history of hypertension (Table 4).

**Table 3.**
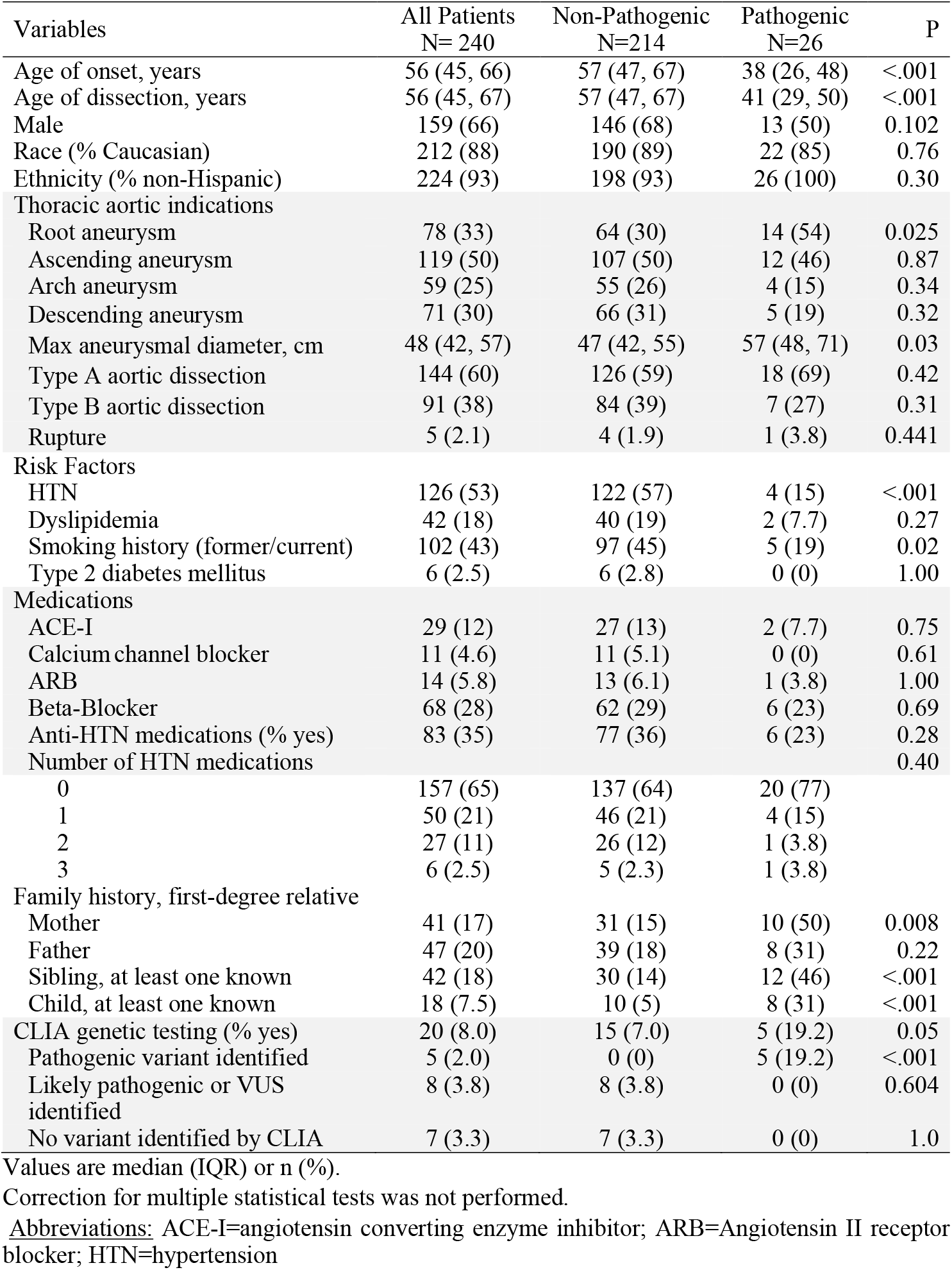
Demographic and Clinical Characteristics at the Time of Dissection

| Variables | All Patients<br>N= 240 | Non-Pathogenic<br>N=214 | Pathogenic<br>N=26 | P |
| --- | --- | --- | --- | --- |
| Age of onset, years | 56 (45, 66) | 57 (47, 67) | 38 (26, 48) | <.001 |
| Age of dissection, years | 56 (45, 67) | 57 (47, 67) | 41 (29, 50) | <.001 |
| Male | 159 (66) | 146 (68) | 13 (50) | 0.102 |
| Race (% Caucasian) | 212 (88) | 190 (89) | 22 (85) | 0.76 |
| Ethnicity (% non-Hispanic) | 224 (93) | 198 (93) | 26 (100) | 0.30 |
| Thoracic aortic indications |  |  |  |  |
| Root aneurysm | 78 (33) | 64 (30) | 14 (54) | 0.025 |
| Ascending aneurysm | 119 (50) | 107 (50) | 12 (46) | 0.87 |
| Arch aneurysm | 59 (25) | 55 (26) | 4 (15) | 0.34 |
| Descending aneurysm | 71 (30) | 66 (31) | 5 (19) | 0.32 |
| Max aneurysmal diameter, cm | 48 (42, 57) | 47 (42, 55) | 57 (48, 71) | 0.03 |
| Type A aortic dissection | 144 (60) | 126 (59) | 18 (69) | 0.42 |
| Type B aortic dissection | 91 (38) | 84 (39) | 7 (27) | 0.31 |
| Rupture | 5 (2.1) | 4 (1.9) | 1 (3.8) | 0.441 |
| Risk Factors |  |  |  |  |
| HTN | 126 (53) | 122 (57) | 4 (15) | <.001 |
| Dyslipidemia | 42 (18) | 40 (19) | 2 (7.7) | 0.27 |
| Smoking history (former/current) | 102 (43) | 97 (45) | 5 (19) | 0.02 |
| Type 2 diabetes mellitus | 6 (2.5) | 6 (2.8) | 0 (0) | 1.00 |
| Medications |  |  |  |  |
| ACE-I | 29 (12) | 27 (13) | 2 (7.7) | 0.75 |
| Calcium channel blocker | 11 (4.6) | 11 (5.1) | 0 (0) | 0.61 |
| ARB | 14 (5.8) | 13 (6.1) | 1 (3.8) | 1.00 |
| Beta-Blocker | 68 (28) | 62 (29) | 6 (23) | 0.69 |
| Anti-HTN medications (% yes) | 83 (35) | 77 (36) | 6 (23) | 0.28 |
| Number of HTN medications |  |  |  | 0.40 |
| 0 | 157 (65) | 137 (64) | 20 (77) |  |
| 1 | 50 (21) | 46 (21) | 4 (15) |  |
| 2 | 27 (11) | 26 (12) | 1 (3.8) |  |
| 3 | 6 (2.5) | 5 (2.3) | 1 (3.8) |  |
| Family history, first-degree relative |  |  |  |  |
| Mother | 41 (17) | 31 (15) | 10 (50) | 0.008 |
| Father | 47 (20) | 39 (18) | 8 (31) | 0.22 |
| Sibling, at least one known | 42 (18) | 30 (14) | 12 (46) | <.001 |
| Child, at least one known | 18 (7.5) | 10 (5) | 8 (31) | <.001 |
| CLIA genetic testing (% yes) | 20 (8.0) | 15 (7.0) | 5 (19.2) | 0.05 |
| Pathogenic variant identified | 5 (2.0) | 0 (0) | 5 (19.2) | <.001 |
| Likely pathogenic or VUS identified | 8 (3.8) | 8 (3.8) | 0 (0) | 0.604 |
| No variant identified by CLIA | 7 (3.3) | 7 (3.3) | 0 (0) | 1.0 |

Values are median (IQR) or n (%).
2 Correction for multiple statistical tests was not performed.
Abbreviations: ACE-I=angiotensin converting enzyme inhibitor; ARB=Angiotensin II receptor blocker; HTN=hypertension

**Table 4.** Risk factors for cases with a pathogenic variant

| Variables | OR | 95% Wald Confidence Limits |  | P-value |
| --- | --- | --- | --- | --- |
|  |  | Lower | Upper |  |
| Age $\leq 50$ vs $> 50$ | 5.5 | 1.6 | 19.7 | 0.008 |
| Sex (female vs male) | 1.1 | 0.3 | 3.8 | 0.84 |
| Caucasian | 0.7 | 0.1 | 3.1 | 0.60 |
| Root aneurysm | 1.7 | 0.6 | 5.2 | 0.34 |
| Hypertension | 5.6 | 1.4 | 22.3 | 0.015 |
| Smoking history | 2.6 | 0.7 | 9.9 | 0.16 |
| Family history |  |  |  |  |
| Mother | 5.7 | 1.4 | 22.3 | 0.013 |
| Father | 0.3 | 0.1 | 1.6 | 0.17 |
| Siblings | 5.1 | 1.1 | 23.9 | 0.04 |
| Children | 6.0 | 1.4 | 26.7 | 0.017 |

Definitions: Hypertension was defined as no hypertension versus had a diagnosis of hypertension. Smoking history was defined as no smoking history versus had a smoking history. Family history was defined as aortic disease noted within a first-degree relative.

## DISCUSSION

The current study reports our initial experience with research-level whole exome sequencing in patients with thoracic aortic dissection or rupture with or without aneurysm. We tested 240 cases and 258 controls for pathogenic variants in 11 genes known to cause aortic dissection^3^. By whole exome sequencing and validation targeted sequencing, we found pathogenic variants in 10.8% of cases and 0% of controls. 58 (27.2%) cases and 51 (19.8%) controls were identified as carriers of variants of unknown significance.

The diagnostic yield of genetic testing of these 11 hereditary thoracic aortic dissection genes in patients with thoracic aortic dissection or rupture could result in a substantial number of patients with improved clinical care. We found a similar yield to that of previous work of identifying pathogenic variants in the same 11 genes (9.3%) based on research exome sequencing of 355 patients with an early age onset (≤56 years) of thoracic aortic dissection who had no family history or syndromic features^14^. In contrast, the yield of a whole exome sequencing in 102 thoracic aortic aneurysm and dissection patients was much lower, only 3.9% of the cases had a pathogenic variants in one of 21 genes of interest^25^. This study^25^ included patients with thoracic aortic aneurysm or thoracic aortic dissection, in contrast to the more severe phenotype of our study population with thoracic aortic dissection and rupture only. A higher incidence of pathogenic variants in the dissection only cases may reflect the more severe presentation in these patients. The 89% of dissection cases that do not have a pathogenic variant may be due to a pathogenic variant currently annotated as a VUS, a pathogenic variant in a gene not yet identified, a high polygenic risk of aortopathy, and/or environmental risk factors. Additional studies of dissection cases may help identify novel genes underlying risk in remaining cases.

In the general population, the incidence of pathogenic variants in our 11 genes of interest is very low (1 × 10^−7^ %, see Supplementary note). However, in our cohort of patients with aortic dissection and rupture, we identified putative disease causing variants in 11 hereditary thoracic aortic dissection genes in 10.8% of cases, which is consistent with another recently published report^3^ and suggests the utility of pursuing a clinical genetic diagnosis in this patient group. The significant risk factors for a pathogenic variant in patients with thoracic aortic dissection or rupture were young age (<50 years old), no history of hypertension, but strong family history of aortic aneurysm, dissection or rupture (especially involving mother, siblings and children) with estimated odds ratios ranging from 5.1 for affected siblings to 6.0 for affected children (Table 4). This is in agreement with a recent study in familial and sporadic cases of aneruysm or dissection of the thoracic aorta which demonstrated a significantly increased probability of harboring a pathogenic or likely pathogenic variant in cases which were syndromic, young (age <50), or with a known or probable family history^26^. Patients with pathogenic variants in *TGFBR1/2* (Loeys-Dietz syndrome), *FBN1* (Marfan syndrome), and *MYH11* have a higher risk of aortic dissection and suffer more complications from aortic dissection, including death^6^. Therefore AHA/ACC guideline recommends early and aggressive prophylactic operation to resect the abnormal thoracic aorta in patients with pathogenic variants.^6^ Based on our results, we would recommend clinical genetic testing of known hereditary thoracic aortic dissection genes for all aortic dissection and rupture patients, especially those with onset < 50 years old, no history of hypertension, and with a family history of aortic dissection. Once a pathogenic variant is identified in the patients, their family members should undergo screening for the same pathogenic variant. If pathogenic variants are identified in any family members, the clinical management will shift towards close monitoring and potentially early prophylactic resection of diseased aorta as recommended by AHA/ACC guidelines.^6^

We did not find a difference in the percentage of variants of unknown significance (VUSs) in 11 dissection genes among cases compared to controls (p = 0.07). In constrast, a previous study found a significantly increased burden of VUSs in hereditary thoracic aortic dissection genes in dissection cases < 56 years of age compared with public controls (p = 2 × 10^−8^).^14^ However, several differences in the two studies may have contributed to these different results. The sample size of the control group was substantially higher in the previous study, however the study presented here had sequenced cases and controls in the same batch and performed all quality control and variant annotation blinded to case or control status. Additionally, a focus on younger onset dissection cases may identify higher rates of VUSs that may actually be pathogenic.^27^ Although the 2015 American College of Medical Genetics guidelines^15^ state that “a variant of uncertain significance should not be used in clinical decision-making” we found evidence that variants of unknown significance from CLIA-certified genetic testing resulted in the introduction of syndromic labels and diagnoses into the medical record. For example, a variant of unknown significance in *TGFBR2* was subsequently described as a “novel change likely causing Loeys-Dietz syndrome.” The statistically similar rate of VUS in cases and controls demonstrates the need for greater understanding of the high frequency of VUSs in controls (15% in Guo et al and 20% in this study) and careful interpretation of variants of unknown significance in clinical practice.

To address the limitation that our sample processing and whole exome sequencing was not CLIA-certified, we verified pathogenic variants using MIPS. Furthermore, we performed expert-annotation of variant pathogenicity blinded to case or control status. This, coupled with the absence of pathogenic variants in controls, provides increased confidence in these results. These precautions lend additional evidence that the research-level whole exome sequencing results are of high enough quality to return findings to patients, which will trigger verification by CLIA-certified genetic testing followed by cascade screening for the same pathogenic variant in family members. An EMR review of the cases with a pathogenic variant suggested an average of 4 (3.88) first degree relatives per patient would now be candidates for cascade screening.

In conclusion, this work provides evidence that whole exome sequencing and annotation can accurately identify pathogenic variants in established genes for hereditary thoracic aortic dissection in patients with a thoracic aortic dissection or rupture. Moreover, the results highlight meaningful implications for precision health by providing clinical guidance on how to manage both patients and family members. We recommend clinical genetic testing of hereditary thoracic aortic dissection genes in patients who have suffered a thoracic aortic dissection, especially those with onset prior to 50 years old, a family history of aortic dissection or rupture, or no history of hypertension. These clinical genetic test results may help to prevent catastrophic events, such as thoracic aortic dissections and death within family members of pathogenic variant carriers who have a high risk, but have not developed thoracic aortic dissection or rupture.

## Supporting information

## ACKNOWLEDGEMENTS

Sequencing/Genotyping services were provided through the RS&G Service by the Northwest Genomics Center at the University of Washington, Department of Genome Sciences, under U.S. Federal Government contract number HHSN268201100037C from the National Heart, Lung, and Blood Institute. The authors acknowledge the University of Michigan Medical School Central Biorepository for providing biospecimen storage, management, and distribution services in support of the research reported in this publication. We acknowledge the University of Michigan DNA Sequencing Core. We thank the clinicians, staff, and study participants from the CHIP Biorepository and Michigan Genomics Initiative.

