## Supplementary material for "Clinical implications of identifying pathogenic variants in aortic dissection patients with whole exome sequencing"

Supplementary Note

**Concordance between research-level whole exome sequencing and CLIA-certified genetic testing**

20 (20/240) aortic dissection cases had previous CLIA-certified genetic testing in their medical record. For 13 patients our findings agreed with CLIA-certified genetic testing (pathogenic=5, no findings=5, VUS=3). The remaining 7 cases had discrepancies between CLIA-certified genetic testing and research-level WES and variant annotation. For one patient we identified a VUS in *MYH11,* which was not one of the 6 genes CLIA tested, and for another patient CLIA identified VUS in 2 genes (*CBS, COL5A1*) which were not in the 11 HTAAD genes we annotated. (Supplementary Table 5).

In another patient, functional annotations and the protein domain affected were sufficient evidence for classification as VUS in both *MYLK* and *COL3A1* which were tested in CLIA-certified genetic testing in 2016 but considered benign. For 1 patient, CLIA-certified genetic testing found double heterozygous genotypes for 2 VUS variants in *COL5A1* and *CBS* which were not annotated in our research-level genetic testing.

3 patients were found to have likely pathogenic or possibly causative variants by CLIA-certified genetic testing which we annotated as VUS. Finally, one patient had a 2012 CLIA-certified genetic testing result of pathogenic which we annotate as a VUS due to lack of evidence for pathogenicity.

**Pathogenic variants in commonly used databases**

Of the 17 of our 24 pathogenic variants present in ClinVar as of September 30, 2018, 12 are pathogenic, 8 are listed as pathogenic, 5 as likely pathogenic, 1 as conflicting interpretation of pathogenicity, and 3 as VUS. 1 of the VUS variants is for a non-aortic phenotype—Wolff-Parkinson-White pattern. 15 of the 24 variants have an rsID in dbSNP v151 for the hg19 chromosomal position, but only 11 of those have reference and alternate alleles corresponding to the variation catalogued in our cohort. For example, dbSNP lists rs113935744 as having reference allele T and alternate allele A whereas our sample was a carrier for alternate allele C.

5,344 loss of function and missense variants from the 11 genes of interest were obtained from gnomAD v2.1. 930 of those variants are listed in ClinVar with the same reference and alternate alleles as gnomAD. 16 of those are Pathogenic or Pathogenic/Likely pathogenic. By summing the allele counts across variants, we estimate pathogenic variants in these genes have a background prevalence of $9.396 \times 10^{-6}$ (30 occurrences in 3,193,956 alleles). Of the 24 pathogenic variants, only 2 are catalogued (rs779512296 and rs761857514) in gnomAD. rs779512296 has an allele frequency of 0.00002891 in the gnomAD Latino population and 0.000008801 in the non-Finnish European population. rs761857514 has an allele frequency of 0.00003267 in the South Asian population.

In this effort we used pathogenicity filtering criteria tailored to our phenotype of interest. As previously shown, using a typical pathogenicity filter (predicted deleterious by at least two of Polyphen2, SIFT, and MutationTaster; 0.5% maximum allele frequency across European Americans and African Americans in the Exome Variant Server; and 5% maximum allele frequency in 1000G) there is a high background prevalence of protein-altering variants in a population[4]. For example, default filtering on GeneVetter (genevetter.kidneyomics.org) identifies 322 of 2,535 (12.7%) 1000 Genomes samples as pathogenic variant carriers, which is a higher background prevalence than we might expect for TAAD. Using the same filter in our cohort, we identify 48 cases (19.3%) and 22 controls (8.5%) as carriers for a pathogenic variant.

**Supplementary Tables**

| **Gene** | **NCBI** |
|---|---|
| *ACTA2* | NM_001141945.1 |
| *COL3A1* | NM_000090.3 |
| *FBN1* | NM_000138.4 |
| *LOX* | NM_002317.5 |
| *MYH11* | NM_002474.2 |
| *MYLK* | NM_053025.3 |
| *PRKG1* | NM_001098512.3 |
| *SMAD3* | NM_005902.3 |
| *TGFB2* | NM_003238.3 |
| *TGFBR1* | NM_004612.2 |
| *TGFBR2* | NM_003242.5 |

**Supplementary Table 1**: mRNA-seq isoforms used to identify the predicted amino acid change. Typically, this is a major isoform expressed in smooth muscle cells. For some proteins, previous publication's isoform was chosen. NM indicates manually annotated and reviewed mRNAs.

| | Pathogenic (N=24) | | | | | | | VUS (N=86) | | | | | | |
|---|---|---|---|---|---|---|---|---|---|---|---|---|---|---|
| **Gene** | **Cases* (N=237)** | **Controls (N=258)** | **Fisher Exact Test p-value** | **Odds Ratio Estimate** | **Odds Ratio 95% Confidence Interval** | **Chi-square test p-value (Yates' continuity correction)** | **Chi-square test statistic (Yates' continuity correction)** | **Cases* (N=213)** | **Controls (N=258)** | **Fisher Exact Test p-value** | **Odds Ratio Estimate** | **Odds Ratio 95% Confidence Interval** | **Chi-square test p-value (Yates' continuity correction)** | **Chi-square test statistic (Yates' continuity correction)** |
| *ACTA2* | NA | NA | | | | | | 1 | 0 | 0.452 | Inf | 0.031, Inf | | |
| *COL3A1* | 1 | 0 | 0.48 | Inf | 0.028, Inf | | | 9 | 5 | | | | 0.237 | 1.4 |
| *FBN1* | 18 | 0 | | | | 2.05e-5 | 18.14 | 12 | 15 | | | | 1 | 1.18e-29 |
| *LOX* | 1 | 0 | 0.48 | Inf | 0.028, Inf | | | NA | NA | | | | | |
| *MYH11* | NA | NA | | | | | | 18 | 13 | | | | 0.194 | 1.69 |
| *MYLK* | NA | NA | | | | | | 5 | 4 | 0.738 | 1.525 | 0.323, 7.789 | | |
| *PRKG1* | 2 | 0 | 0.23 | Inf | 0.204, Inf | | | 3 | 3 | 1 | 1.213 | 0.161, 9.16 | | |
| *SMAD3* | 2 | 0 | 0.23 | Inf | 0.204, Inf | | | 4 | 0 | 0.041 | Inf | 0.805, Inf | | |
| *TGFB2* | NA | NA | | | | | | 5 | 3 | 0.477 | 2.040 | 0.392, 13.290 | | |
| *TGFBR1* | NA | NA | | | | | | 3 | 3 | 1 | 1.214 | 0.161, 9.16 | | |
| *TGFBR2* | 1 | 0 | 0.48 | Inf | 0.028, Inf | | | 7 | 9 | | | | 1 | 7e-31 |

Supplementary Table 2. Association between variants of a given class and case/control status per each of the 11 HTAAD genes. A sample from each of the two related pairs in the cases was removed while the first ascertained sample was retained. When testing the VUS class of variants, only cases without a pathogenic variant were considered. Accounting for multiple testing using a Bonferroni threshold of 0.003, the only significant association identified is for pathogenic variants in *FBN1*.

| Chr | Pos | Variant type | Ref | Alt | Sample (NHLBI_ID) | Sample (GWAS/MIPS ID) | WES (GT:AD:DP:GQ:PL) | MIPS_v1 Variant call (GT:DP:GQ:PL) | MIPS_v1 Quality | MIPS_v2 Variant call (GT:DP:GQ:PL for SNPs, GT:PL:DP:AD:GQ for indels) | MIPS_v2 Quality |
|---|---|---|---|---|---|---|---|---|---|---|---|
| 15 | 48707913 | SNP | T | C | 16554 | 58432 | 0/1:28,17:45:99:488,0,896 | 0/1:267:99:255,0,255 | Failed individual level call rate filter | 0/1:676:99:255,0,255 | Pass |
| 15 | 48713849 | SNP | G | C | 19082 | 113392 | 0/1:36,35:71:99:952,0,1142 | NA | Sample not sequenced | 0/1:222:255:255,0,255 | sample filtered out due to high missingness in first pass, variant filtered by SVM filter |
| 15 | 48714232 | SNP | C | A | 11353 | 57411 | 0/1:43,33:76:99:931,0,1329 | 0/1:1165:99:255,0,255 | Pass | 0/1:1050:99:255,0,255 | Pass |
| 15 | 48719947 | Indel | TGAAGCAGTACCCTTCCC | T | 17339 | 57403 | 0/1:26,12:38:99:427,0,4465 | NA | Indel calling not performed | 0/1:1189:.:583,586,20:43177,0,38932 | Pass |
| 15 | 48722967 | SNP | A | G | 15731 | 58466 | 0/1:8,6:14:99:175,0,237 | 0/1:1065:99:255,0,255 | Pass | 0/1:742:99:255,0,255 | Pass |
| 15 | 48725107 | SNP | C | T | 12040 | 57445 | 0/1:19,16:35:99:427,0,631 | 0/1:607:99:255,0,255 | Failed individual level call rate filter | 0/1:1564:99:255,0,255 | Pass |
| 15 | 48730109 | SNP | G | A | 11487 | 57396 | 0/1:12,7:19:99:216,0,401 | 0/1:86:99:255,0,255 | Pass | 0/1:162:99:255,0,255 | Pass |
| 15 | 48744873 | SNP | C | T | 16426 | 113380 | 0/1:13,13:26:99:318,0,361 | NA | Sample not sequenced | 0/1:242:99:255,0,255 | Pass |
| 15 | 48760660 | SNP | A | G | 17258 | 113401 | 0/1:28,35:63:99:975,0,807 | NA | Sample not sequenced | 0/1:154:255:255,0,255 | sample filtered out in second pass due to missingness rate, variant passes filter |
| 15 | 48764793 | SNP | A | G | 15339 | 57412 | 0/1:22,24:46:99:724,0,693 | 0/1:3287:99:255,0,255 | Pass | 0/1:8893:99:255,0,255 | Pass |
| 15 | 48773870 | Indel | C | CT | 12144 | 57419 | 0/1:24,29:53:99:741,0,561 | NA | Indel calling not performed | 0/1:1748:.:836,905,7:24198,0,21469 | Pass |
| 15 | 48782066 | SNP | C | A | 11080 | 58472 | 0/1:32,20:52:99:533,0,1034 | 0/1:881:99:255,0,255 | Failed individual level call rate filter | 0/1:2433:99:255,0,255 | Pass |
| 15 | 48786401 | SNP | C | G | 16641 | 57386 | 0/1:49,52:101:99:1347,0,1369 | 0/1:104:99:255,0,255 | Pass | 0/1:218:99:255,0,255 | Pass |
| 15 | 48802264 | Indel | G | GT | 11970 | 57402 | 0/1:27,38:65:99:1212,0,802 | NA | Indel calling not performed | 0/1:2213,0,2047:162:79,83,0:. | sample filtered out due to high missingness in first pass |
| 15 | 48802366 | SNP | T | C | 15837 | 57597 | 0/1:12,16:28:99:460,0,355 | 0/1:372:99:255,0,255 | Pass | 0/1:240:99:255,0,255 | Pass |
| 15 | 48808561 | SNP | T | C | 13555 | 113354 | 0/1:19,16:35:99:525,0,561 | NA | Sample not sequenced | 0/1:415:99:255,0,255 | Pass |
| 15 | 48812913 | SNP | G | A | 16930 | 57832 | 0/1:27,26:53:99:773,0,755 | 0/1:427:99:255,0,255 | Pass | 0/1:753:99:255,0,255 | Pass |
| 15 | 48888576 | SNP | C | T | 17920 | 57617 | 0/1:21,16:37:99:480,0,699 | NA | No coverage in sequencing bam file | 0/1:525:99:255,0,255 | Pass |
| 15 | 67457370 | Indel | TGAA | T | 10317 | 57605 | 0/1:17,19:36:99:727,0,647 | NA | Indel calling not performed | 0/1:180:.:95,85,0:2617,0,3122 | Pass |
| 15 | 67462935 | Indel | TA | T | 16115 | 58000 | 0/1:25,41:66:99:1336,0,736 | NA | Indel calling not performed | 0/1:89:.:49,39,1:1005,0,1307 | Pass |
| 15 | 67462935 | Indel | TA | T | 13332 | 58351 | 0/1:40,27:67:99:810,0,1268 | NA | Indel calling not performed | 0/1:39:.:16,23,0:632,0,396 | Pass |

| | | | | | | | | | | | |
|---|---|---|---|---|---|---|---|---|---|---|---|
| 2 | 189858169 | SNP | G | A | 15202 | 57577 | 0/1:46,36:82:99:1077,0,1385 | 0/1:363:99:255,0,255 | Pass | 0/1:446:99:255,0,255 | Pass |
| 3 | 30732950 | SNP | G | A | 17845 | 57458 | 0/1:16,23:39:99:667,0,499 | 0/1:234:99:255,0,255 | Pass | 0/1:621:99:255,0,255 | Pass |
| 10 | 53227579 | SNP | G | A | 19825 | 57370 | 0/1:42,47:89:99:1590,0,1192 | 0/1:39:99:255,0,255 | Failed individual level call rate filter | 0/1:301:99:255,0,255 | Pass |
| 10 | 53227579 | SNP | G | A | 10712 | 57607 | 0/1:91,58:149:99:1689,0,2706 | 0/1:2002:99:255,0,255 | Pass | 0/1:168:255:255,0,255 | sample filtered out in second pass due to high missingness |
| 5 | 121412592 | Indel | CCAGA | C | 14301 | 57653 | 0/1:44,39:83:99:1506,0,2525 | NA | Indel calling not performed | 0/1:1650:.:757,879,14:31292,0,25368 | Pass |

**Supplementary Table 3. Confirmation of WES variant calls with Molecular Inversion Probe Sequencing (MIPS)**. Two rounds of MIPS were performed to confirm the pathogenic variant calls in all 26 patients. In round 1, 22 of the 26 samples were sequenced. In round 2, all samples were sequenced.

| | CLIA | | | Research | | | |
|---|---|---|---|---|---|---|---|
| **Sample** | **CLIA year** | **Clinical Genetic Results from EMR** | **Annotation** | **Variant** | **Annotation** | **Gene** | **Rationale for discrepancy** |
| 1 | 2015 | Heterozygous for the p.R192Q pathogenic mutation in the *PRKG1* gene | Pathogenic | 10:53227579 | Pathogenic | *PRKG1* | Concordant |
| 2 | 2010 | Mutation: *FBN1* Exon 22 Nucleotide: c.2728G>C Amino Acid:Asp910His | Pathogenic | 15:48786401 | Pathogenic | *FBN1* | Concordant |
| 3 | NA | Genetically confirmed Marfan Syndrome | Pathogenic | 15:48782066 | Pathogenic | *FBN1* | Concordant |
| | | | | 2:189856434 | VUS | *COL3A1* | Concordant |
| 4 | NA | Clinical genetic testing, no variant identified | No findings | NA | NA | *NA* | Concordant |
| 5 | 2014 | Panel was negative for everything. *COL3A1 TGFBR1 TGFBR2, ACTA2, SMAD3, TGFB2* tested. | No findings | 16:15820794 | VUS | *MYH11* | *MYH11* not tested in CLIA panel |
| 6 | 2012 | *SMAD3* genetic mutation | Pathogenic | 15:67462935 | Pathogenic | *SMAD3* | Concordant |
| 7 | 2012 | VUS from *TGFBR2* | VUS | 3:30713866 | VUS | *TGFBR2* | Concordant |
| 8 | 2016 | No genetic mutations discovered, 22 gene panel including *COL3A1* and *MYLK* | No findings | 3:123337545 | VUS | *MYLK* | *MYLK* p.T1814I is absent in the ExAC and gnomAD database. T1814 alteration is not reported before so it is unclear whether alter this amino acid lead to TAD. Multiple functional prediction programs suggest that this variant is damaging. Classified as VUS based on ACMG guidelines. |
| | | | | 2:189863424 | VUS | *COL3A1* | Variant is in triple helical region but didn't alter critical glycine |
| 9 | NA | 6 gene vascular aneurysm panel and fibrillin 1 sequencing were negative | No findings | NA | NA | NA | Concordant |
| 10 | NA | *SMAD3* mutation related to Loeys-Dietz syndrome | Pathogenic | 15:67462935 | Pathogenic | *SMAD3* | Concordant |
| 11 | 2017 | Patient was negative for panel | No findings | NA | NA | NA | Concordant |
| 12 | 2014 | *SMAD3* likely pathogenic variant | Likely pathogenic | 16:15844048 | VUS | *MYH11* | *MYH11* p.K1256del is not found in the ExAC and gnomAD database. Deletion of this amino acid is not reported before so it is unclear whether deletion of this amino acid leads to disease. A couple of single amino acid deletions flanking K1256 are found in the gnomAD and ExAC datebases. In the gnomAD v2.1 control database, there are 6 K1263del alleles and 2K1231del alleles. Classified as VUS based on ACMG guidelines. |
| | | | | 15:67482824 | VUS | *SMAD3* | *SMAD3* p.V410 is found in the ExAC with low MAF (5.53E-04). T1814 alteration is not reported before so it is unclear whether altering this amino acid leads to disease. Some |

| | | | | | | | |
|---|---|---|---|---|---|---|---|
| | | | | | | | functional prediction programs suggest that this variant is damaging and others suggest benign. Classified as VUS based on ACMG guidelines. |
| 13 | 2014 | *SMAD3* gene mutation in exon 9, c.1228G>T, p.Val410Phe | Likely pathogenic | 15:67482824 | VUS | *SMAD3* | Same as above |
| 14 | NA | Only was tested for Marfan and was found to be negative | No findings | NA | NA | NA | Concordant |
| 15 | 2012 | Possibly causative *SMAD3* mutation (c.331T>A) | Possibly causative | 15:67457357 | VUS | *SMAD3* | Lack evidence for pathogenicity, therefore classified as VUS based on ACMG guidelines. |
| 16 | 2016 | VUS in *COL3A1* p. V5291 | VUS | 2:189860493 | VUS | *COL3A1* | Concordant |
| 17 | 2016 | Heterozygous for the p.R369C pathogenic mutation in the *CBS* gene. Heterozygous for the p.P435A (c.1303C>G) VUS in the *COL5A1* gene | VUS | 21:44480591 | NA | *CBS* | Not one of 11 HTAAD genes |
| | | | | 9:137623480 | NA | *COL5A1* | Not one of 11 HTAAD genes |
| 18 | 2012 | *FBNI* exon 32 Nucleotide: c. 4057G>A Amino: Gly1353Arg | Likely pathogenic | 15:48766755 | VUS | *FBN1* | Reported in patients, no evidence for pathogenicity. Located in EGF-like 22 calcium binding domain and is not a critical amino acid for the domain. |
| 19 | 2013 | *TGFBR1* Exon 5 Nuc: c.949C>T AA: His317Tyr | Likely pathogenic | 9:101904961 | VUS | *TGFBR1* | Lack evidence for pathogenicity, therefore classified as VUS based on ACMG guideline. |
| 20 | 2013 | No mutations found | No findings | NA | NA | NA | NA |

Supplementary Table 4. Concordance between research-level and CLIA-certified genetic testing in the 20 patients with CLIA genetic testing results in our cases. The variants calls were always concordant, the discrepancies were in interpretation of evidence for annotation as pathogenic or VUS.

| **Variant class (# of variants in class)** | | **Cases** | **Controls** | **Chi-square test p-value (Yates' continuity correction)** | **Chi-square test statistics (Yates' continuity correction)** |
|---|---|---|---|---|---|
| | | **n=238** | **n =258** | | |
| **pathogenic (24)** | Non-carrier | 213 | 258 | 2.79e-7 | 26.39 |
| | Carrier | 25 | 0 | | |
| | | **n=213** | **n=258** | | |
| **VUS (86)** | Non-carrier | 155 | 207 | 0.072 | 3.25 |
| | Carrier | 58 | 51 | | |

Supplementary Table 5. Association between variants of a given class and case/control status across all 11 genes. A sample from each of the two related pairs in the cases was removed while the first ascertained sample was retained. When testing the VUS class of variants, only cases without a pathogenic variant were considered.
